# Fluid-Structure Interactions of Peripheral Arteries Using a Coupled *in silico* and *in vitro* Approach

**DOI:** 10.1101/2023.04.17.537257

**Authors:** S. Schoenborn, T. Lorenz, K. Kuo, D.F. Fletcher, M. A. Woodruff, S. Pirola, M. C. Allenby

## Abstract

Vascular compliance is considered both a cause and a consequence of cardiovascular disease and a significant factor in the mid- and long-term patency of vascular grafts. However, the biomechanical effects of localised changes in compliance, such as during plaque development or after bypass grafting and stenting, cannot be satisfactorily studied with the available medical imaging technologies or surgical simulation materials. To address this unmet need, we developed a coupled *silico-vitro* platform which allows for the validation of numerical fluid-structure interaction (FSI) results as a numerical model and physical prototype. This numerical one-way and two-way FSI study is based on a three-dimensional computer model of an idealised femoral artery which is validated against patient measurements derived from the literature. The numerical results are then compared with experimental values collected from compliant arterial phantoms. Phantoms within a compliance range of 1.4 - 68.0%/100mmHg were fabricated *via* additive manufacturing and silicone casting, then mechanically characterised *via* ring tensile testing and optical analysis under direct pressurisation with differences in measured compliance ranging between 10 - 20% for the two methods. One-way FSI coupling underestimated arterial wall compliance by up to 14.71% compared to two-way FSI modelling. Overall, Smooth-On Solaris matched the compliance range of the numerical and *in vivo* patient models most closely out of the tested silicone materials. Our approach is promising for vascular applications where mechanical compliance is especially important, such as the study of diseases which commonly affect arterial wall stiffness, such as atherosclerosis, and the model-based design, surgical training, and optimisation of vascular prostheses.

## 1 Introduction

Cardiovascular disease (CVD) is the most common cause of death worldwide. The annual incidence of global deaths related to CVD is expected to rise from 16.7 million in 2002 to 23.3 million in 2030 [1].

A common manifestation of CVD is peripheral artery disease (PAD), whose symptoms develop as a consequence of the critical narrowing of peripheral arteries. International research studies show a prevalence of approximately 10 - 20% for PAD depending on study design, country, and participant’s age and gender [2]. This paper focuses on the femoropopliteal artery since it is the most likely segment of arterial tissue to develop PAD [3, 4].

Two of the main causes of PAD are atherosclerosis, which is a chronic and progressive vascular disease characterised by the inflammation of the tunica intima and the accumulation of lipoproteins in the stiffening arterial walls [5], and intimal hyperplasia (IH), the chronic and excessive thickening of the tunica intima, caused by abnormal proliferation and migration of vascular smooth muscle cells (VSMCs) in response to endothelial injury or dysfunction with associated deposition of connective extracellular matrix [6-8]. Both atherosclerosis and IH can cause significant narrowing of affected blood vessels, called stenosis [9], potentially leading to claudication or critical limb ischemia [10]. While the exact causes of atherosclerosis are unknown, there are many risk factors which have been shown to increase the likelihood of developing this disease. These factors include lifestyle and biological factors such as high total cholesterol, diabetes, obesity, hypertension, old age, stress, lack of exercise, and poor diet, among others [11, 12]. Additionally, vascular disease initiation or progression, such as atherosclerosis and IH in a vessel, are often related to adverse vascular hemodynamics and biomechanics [13].

One of the most important biomechanical signals affecting the phenotypes of endothelial cells (ECs) and VSMCS is wall shear stress (WSS). Non-physiological or oscillatory WSS may lead to EC misalignment and inflammation or cause VSMCs to change their phenotype to a proliferative state, which causes them to proliferate and migrate towards the tunica intima where they accumulate and deposit excessive extracellular material [14, 15]. Both non-physiologically high and low WSS have been reported to affect the development and progression of atherosclerosis and IH, where the physiological range depends on vessel location and composition [16]. Another biomechanical signal playing a significant role in vascular diseases is intramural stress or wall stress (WS). High WS may serve as an initiator of the EC inflammatory response and injury, which is a promoting factor for VSMC proliferation [17]. WS and WSS are directly related to local hemodynamics and arterial wall properties such as mechanical compliance [18], with adverse biomechanics often occurring following disease or vascular interventions [19, 20].

The pathophysiological behaviour of arteries can be analysed *in vivo, in vitro*, and *in silico. In vivo* analyses using medical imaging technologies are increasing in popularity. However, relevant flow parameters, such as wall shear stress and intramural stress in a blood vessel, cannot be directly measured *in vivo* [21]. While 4D magnetic resonance imaging (MRI) and phase contrast MRI have been used for the validation of vascular simulations, their use for estimating WSS through spatial derivation of a velocity field measured in clinical practice remains limited due to poor spatial resolution along the arterial wall [22-25]. To capture relevant velocity information using 4D MRI, it is recommended to measure at least 5 - 6 voxels across the vessel diameter of interest with typical spatial resolutions between 1.5×1.5×1.5 and 3×3×3 mm^3^ [26], which limits use in small-diameter applications. Therefore, physical (*in vitro*) and computational (*in silico*) models remain essential to support the design of vascular prostheses and pre-operative planning.

Numerical modelling is considered a valid alternative to clinical studies for quantitative hemodynamic research [27] and aids the study of physical phenomena that cannot be easily measured *in vivo*. The use of fluid-structure interaction (FSI) methods enables the coupling of fluid and structural domains to study complex phenomena between the blood flow and deformable arterial wall. While FSI simulations could potentially generate more accurate results for WSS and WS [28, 29], the addition of a compliant vascular wall poses new challenges for numerical modelling and its experimental validation. In systems with large deformations such as blood flow in arteries, a computationally more expensive two-way coupling is usually required [30] and is becoming more standard as sufficient computational power is becoming more accessible [31-35]. However, some studies on peripheral arteries apply one-way coupling with the argument that the deformation of the arterial wall in these locations is negligible [36-39].

Experimental models utilising an arterial phantom are valuable for patient-specific research experiments [40], vascular device testing [41], surgical training [42], and validation of numerical model results [43], but many fail to mimic physiological arterial compliance which has been shown to reduce the precision of the model due to overestimation of WSS and underestimation of recirculation zones [44], which are risk factors for atherosclerosis and IH [13]. Various types of materials have been described in the literature for the manufacturing of compliant arterial phantoms, which include flexible resins such as Formlabs Flexible and Stratasys Polyjet materials, such as FullCure 930 TangoPlus [45, 46]. Polymethyldisiloxane (PDMS) is a frequently-used model material which is highly optically transparent (75 - 92% transmittance [47]) but care must be taken to ensure defined curing temperatures [48] and mixing ratios [49] during arterial phantom manufacturing due to their significant effect on the mechanical properties of the cured silicone.

The goal of this paper is to fill this gap by proposing a coupled numerical and experimental workflow to create compliant *in silico* and *in vitro* arterial and silicone models and validate their mechanical properties against published patient data. In this paper, idealised femoral arterial phantoms will be manufactured from Sylgard 184 (PDMS) and two other commercially available platinum-cured silicones, Solaris™ and PlatSil® Gel-OO20, *via* additive manufacturing and silicone moulding. These experimental models will be assessed on their ability to replicate physiological vascular compliance using both direct internal pressurisation and ring tensile testing including a comparison of the two methods. Lastly, one-way and two-way coupling in peripheral vascular FSI models will be compared for spatial and temporal differences between generated results in both the structural and fluid domain.

## 2 Methodology

### 2.1 Experimental Methods

#### 2.1.1 Manufacturing of Silicone Samples

In this experimental study, small-diameter arterial phantoms mimicking idealised femoral arteries with wall thicknesses of 0.8 mm, 1.2 mm, and 1.6 mm [50] were created *via* additive manufacturing and casting of three different silicones (Figure 1A). The models were assessed on the basis of circumferential compliance, which was derived from optical measurements under direct pulsatile pressurisation and from uniaxial ring tensile test data.

**Figure 1:**
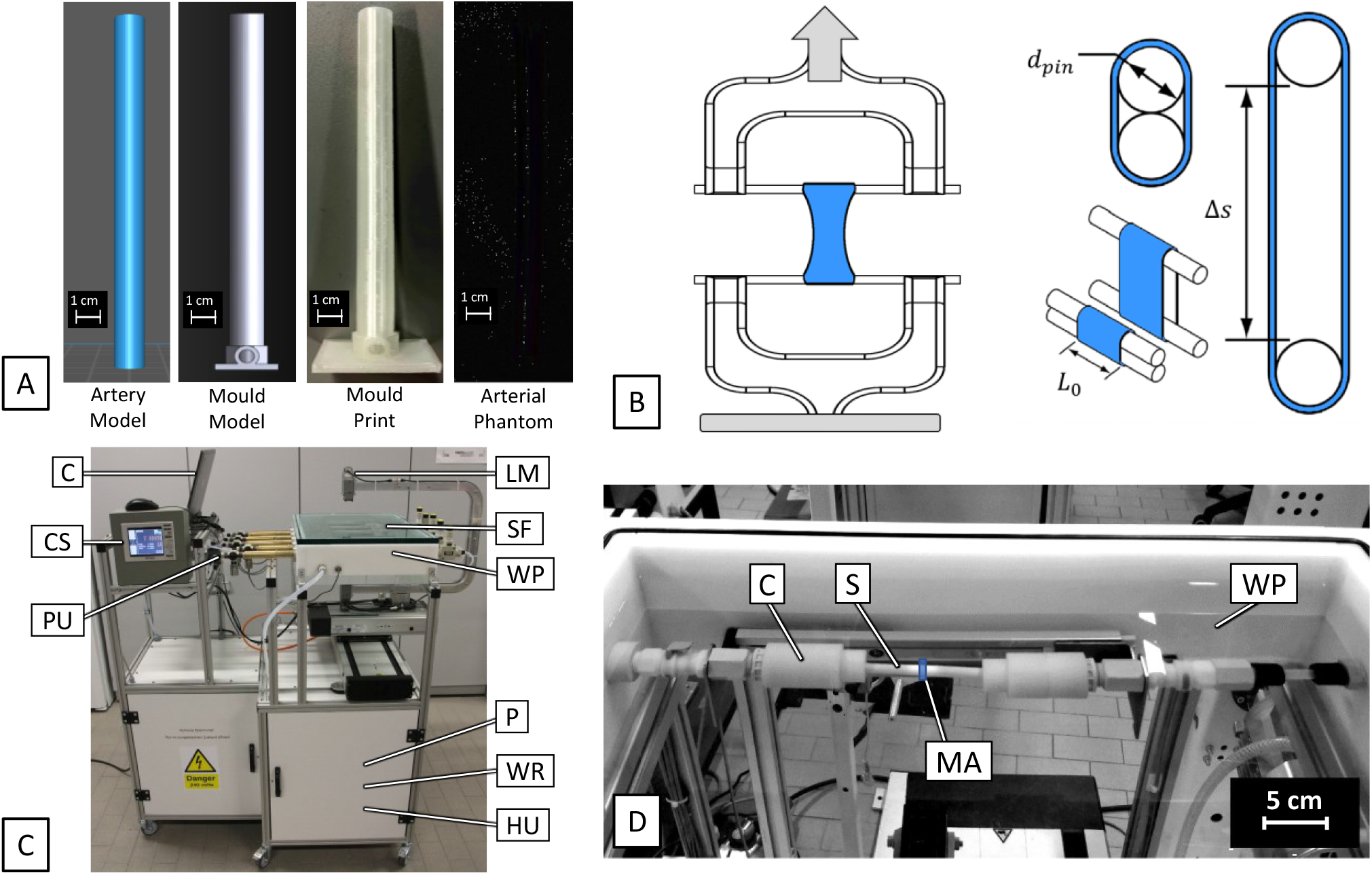
Manufacturing and testing of idealised arterial phantoms. A) CAD modelling, 3D printing, and moulding process of silicone arterial phantoms; B) Schematic of ring tensile testing of arterial phantoms with d_pin_: pin diameter, L_0_: initial sample length, Δs: displacement; C) Vascular testing rig for pulsatile direct pressurisation of arterial phantoms. C: Computer, CS: Control System, PS: Pulsation Unit, LM: Laser Micrometer, SF: Sample Fixation, WP: Water Pool, P: Pump, WR: Water Reservoir, HU: Heating Unit; D) View of the arterial phantom in the pulsatile direct pressurisation rig. C: Clamp, S: Sample, MA: Measurement Area, WP: Water Pool.

The moulds used for silicone casting were 3D printed on a Raise3D Pro2 Plus (Raise3D) fused deposition modelling (FDM) printer with a 0.4 mm nozzle using Raise3D Premium Polyvinyl Alcohol (PVA) 1.75 mm diameter filament (Raise3D). Every mould was printed with a 0.2 mm layer height using 3 perimeter shells and 100% infill density. “Stringing”, a phenomenon commonly occurring with the additive manufacturing of PVA [51] where excess material oozes from the nozzle without extruder movement, was minimised to avoid holes in the final phantom. This was achieved by storing the filament in an eSUN eBox Filament Storage Dry Box heated to 45°C with a small bag of desiccant to prevent absorption of moisture, applying no 3D printer collection plate heating, and applying a low 3D printer nozzle temperature of 200°C.

The silicone materials compared in this study are Solaris™ (Smooth-On), Sylgard® 184 (Dow Corning), and PlatSil® Gel-OO20 (Polytek), which are chemically curing two-component products. To prepare the silicones, the two components were drawn separately into two syringes, added to a cup according to the volume ratio recommended by the respective manufacturer, and manually stirred for 3 minutes to ensure sufficient mixing.

The mixture was degassed before it was injected into the mould in order to remove any bubbles that were introduced into the silicone gel during stirring and prevent holes in the final model. To facilitate this, the silicone gel was placed into an airtight container connected to a RS-1 / 2.5CFM Rotary Vane Mini Single Stage Vacuum Pump (Taizhou Kooya Vacuum Technology) and exposed to a gauge pressure of -0.8 bar for 20 minutes. During this process, the silicone gel foamed up as air bubbles were pulled to the surface. At the end of the degassing process, no more bubbles were visible. Air was then slowly reintroduced until the pressure levels returned to atmospheric pressure and the cup with the silicone gel was removed from the vacuum chamber.

The mixed and degassed silicone gel was carefully drawn into a luer slip syringe. During this process, care was taken to not re-introduce air bubbles into the system. The silicone was then slowly injected into the appropriately sized sprues at the bottom of the moulds until it reached the open top of the mould. The sprues were then closed with small plugs that have been prepared via FDM printing to prevent the silicone left from leaking out. The silicone was left to cure inside the moulds at room temperature for 24 hours.

After the curing process was completed, the moulds were fully immersed in a container filled with room-temperature water until the PVA was completely dissolved. Once the arterial phantoms were completely demoulded and free of PVA, the ends of the phantoms were post-processed by cutting off the sprues. Mechanical testing (as described in Sections 2.1.2 and 2.1.3) was conducted no earlier than 14 days after demoulding, since it may take several days for the silicone to achieve its final mechanical properties [52]. Wall thickness and internal diameter were then assessed via light microscopy on a Mikroskops DM4000DM (Leica Microsystems). Additionally, silicone dog bone tensile samples (ASTM D412-C) were manufactured following the same procedure to inform the numerical silicone model.

#### 2.1.2 Tensile Testing

Ring tensile testing is a mechanical testing method that provides accurate information about the behaviour of a sample under internal radial loads and has been reported to yield comparable results with methods utilising direct pressurisation [53]. The ring tensile testing method consists in inserting two small pins into the lumen of the tubular sample. One pin is fixed while the other one is moved at a constant strain rate to increase the distance between the pins until the sample fails (Figure 1B).

A Z2.5 tensile testing machine (Zwick Roell Group) was used to control pin movement. Two adapters were used to provide a rigid support at both ends of the pins (*d*_*pin*_ = 1.5 mm) and allow them to be inserted into the clamps of the tensile testing machine while also allowing them to touch one another. The system provided measurement of pin displacement through its control system and force was registered using a 200 N Xforce HP load cell (Zwick Roell Group). Samples (n = 5 per group) of 10 mm in length were extended until failure at a constant rate of 171 mm/min, mimicking a strain rate experienced by a femoral artery when distending under pressures of 80 to 120 mmHg with 73 bpm [54].

Circumferential compliance was calculated using the elasticity theory as provided by Timoshenko [55, 56], in units of %/100mmHg:

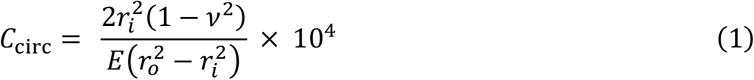

using internal radius *r*_*i*_, external radius *r*_*0*_, Poisson’s ratio *v* and Young’s modulus *E*. The Poisson’s ratio is assumed to be *v* = 0.5 [57] and the Young’s modulus *E* for the region of interest between 80 mmHg and 120 mmHg is calculated as:

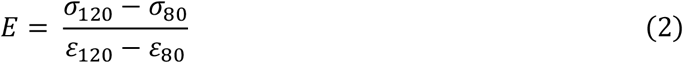

using stress *σ* and strain *ε*. The associated tensile forces *F* were calculated by equating hoop stress *σ*_*h*_ according to Barlow’s formula [58] and the tensile stress within the sample wall *σ*_*t*_ for a given pressure *P* and wall thickness *t*_*0*_ [59]:

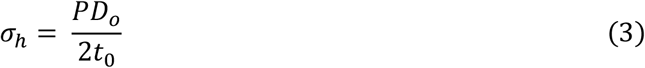

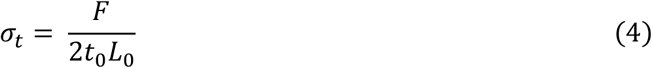

The equivalent external diameter *D*_*o*_ for any extension *s* and initial distance of the outside of the pins *s*_*0*_ was calculated from the perimeter of the sample *U*_*o*_ as follows:

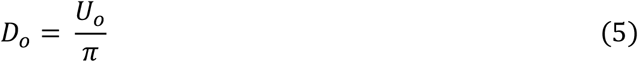

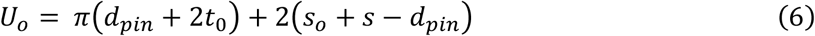

for a given pin diameter *d*_*pin*,_ initial pin distance *s*_*0*_, and displacement *s*.

The material models for the numerical simulations of the silicone behaviour were derived from uniaxial tensile tests of ASTM D412-C dog bone samples utilising the same machine, methodology, and strain rate.

#### 2.1.3 Direct Pressurisation

For dynamic compliance measurements, a pulsatile intraluminal sinusoid pressure profile of 120/80 mmHg was applied using an IWAKI MD-55R Magnetic Drive Pump (IWAKI Singapore Pte Ltd., Singapore) within a closed perfusion loop. The average volume flow of 70 ml/min had a pulse frequency of 69/min and was heated to 35°-39°C. All samples (n = 5 per group) were at least 80 mm in length and the outer diameters of the samples were continuously measured with a LS 7030 LED/CCD laser micrometre (KEYENCE Corporation) at the centre of each sample to avoid entrance and exit effects. The intraluminal pressure was monitored at the sample outlet using a PA3509 pressure gauge with ceramic measuring cell (ifm electronic gmbh). Both signals were recorded and processed using LabVIEW® (National Instruments Austin). Circumferential compliance C_circ_ was calculated from the measurements of pressure P and a calculated inner diameter D_i_ using the following equation [60-62], in units of %/100mmHg:

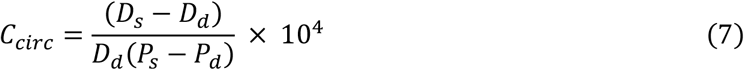

where *P* is pressure (mm Hg), *D* is diameter (mm), and subscripts *s* and *d* are the systolic and diastolic conditions, respectively.

### 2.2 Numerical Methods

An FSI model coupling blood flow and wall motion was developed using the 2022 R1 version of the commercial computational software suite ANSYS (ANSYS Inc., Pennsylvania). ANSYS Mechanical was used for the structural mechanics analysis and ANSYS CFX was used for the fluid mechanics computation. ANSYS Mechanical is a finite-element-based solver for structural mechanics analysis. ANSYS CFX is a finite-volume-based solver with vertex-centred numeric. Three-dimensional continuity and Navier-Stokes equations are used as the governing equations of blood flow [35].

#### 2.2.1 Structural Model

##### 2.2.2.1 Material Models

The femoral arterial models were approximated as cylindrical tubes which match the cross-sectional areas of reference patients aged 25, 45, and 65 years at diastolic pressure as described in the Pulse Wave Database (PWD) published by Charlton et al. [54]. An initial length of L_0_ = 100 mm was selected to allow for sufficient flow development in a transient analysis, the initial total wall thicknesses *t*_*0*_ of 0.8, 1.2, and 1.6 mm were taken from measurements on healthy subjects in the literature [63].

A hyperelastic first-order Ogden model was utilised to reproduce the arterial wall’s stress-strain behaviour [64-66]. The Ogden model parameters were determined using the ANSYS hyperelastic curve fitting function on circumferential uniaxial tensile testing data of healthy femoral arteries reported in literature [67, 68]. The first-order Ogden model is defined as follows [69]:

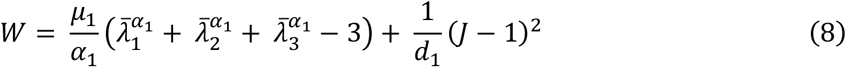

where *W* is the strain energy potential, 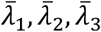 are the deviatoric principal stresses, defined as 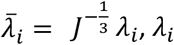 are the principal stresses of the left Cauchy-Green tensor, *J* is the determinant of the elastic deformation gradient, and *μ*_*i*_, *α*_*i*_ and *d*_*k*_ are material constants.

The initial shear modulus μ_0_ is defined by:

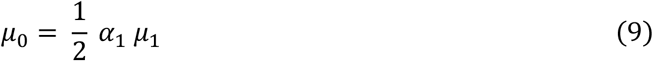

and the initial bulk modulus *K*_0_ is defined by:

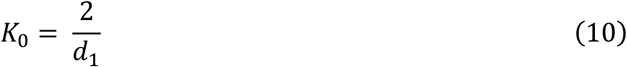

The model parameters utilised for the simulated patient ages without PAD are summarised in Table 1. The tissue surrounding the femoral artery was expected to have a damping effect on the arterial wall movement, which was taken into account by applying a linear elastic support boundary condition [70]. To facilitate a nodal comparison in structural parameters between the two-way and one-way FSI models, the same structural mesh was utilised for all simulation of a given age or wall thickness. The structural model was run as a transient analysis using the sparse direct solver with a time-step size of 10^−3^ s. A timestep was considered as converged when the force convergence value and the displacement convergence value were both below a residual target of 10^−4^.

**Table 1:** First-order Ogden model parameters of structural arterial wall model. Incompressibility parameter d_1_, material constant α_1_, material constants μ_1_ for different patient ages, density ρ.

| Parameter | Value | Source [Reference] |
| --- | --- | --- |
| $\mu_{1,25}$ | 10.100 kPa | Ansys curve fitting tool, source data from literature [71] |
| $\mu_{1,45}$ | 16.625 kPa | Ansys curve fitting tool, source data from literature [71] |
| $\mu_{1,65}$ | 26.414 kPa | Ansys curve fitting tool, source data from literature [71] |
| $\alpha_1$ | 14.405 | Ansys curve fitting tool, source data from literature [71] |
| $d_1$ | $3.63 \times 10^{-6} \text{ Pa}^{-1}$ | Literature [64] |
| $\rho$ | $1080 \frac{\text{kg}}{\text{m}^3}$ | Literature [72] |

The silicone arterial phantom model was approximated as a cylindrical tube which matches the measured diameter and thickness of the average manufactured arterial phantom for the 0.8 mm, 1.2 mm, and 1.6 mm wall thickness groups. The mechanical behaviour of the silicone models was modelled as an isotropic linear elastic material with elastic modulus measured *via* uniaxial tensile testing and with Poisson’s ratio and density informed by literature (Table 2). All silicones were assumed to be nearly incompressible [73].

**Table 2:** Isotropic linear elastic model parameters of structural silicone model. Young’s Modulus E, Poisson’s Ratio v, density ρ.

| Parameter | PDMS | PlatSil | Solaris |
| --- | --- | --- | --- |
| $E$ | $1.655 \pm 0.223 \text{ MPa}$ | $0.205 \pm 0.036 \text{ MPa}$ | $0.539 \pm 0.128 \text{ MPa}$ |
| $\nu$ | 0.499 [74] | 0.499 | 0.499 |
| $\rho$ | $1026.91 \frac{\text{kg}}{\text{m}^3}$ [75] | $1046.85 \frac{\text{kg}}{\text{m}^3}$ [76] | $987.03 \frac{\text{kg}}{\text{m}^3}$ [77] |

##### 2.2.1.2 Iterative Pre-Stress Pipeline

When vascular patient imaging data are collected from *in vivo* medical imaging methods, reconstructed three-dimensional geometries represent a configuration already under load. The most significant loads on the arterial wall are blood pressure, pressure from the surrounding tissue, and residual stresses within the arterial wall itself [78]. Assuming a geometric configuration generated from patient image data to be stress-free, and thus neglecting pre-stress, leads to non-physiologically large deformations when blood pressure is applied to the model during simulation. An inverse design approach, i.e., creating a configuration that will match the imaged geometry when relevant stresses are applied, has been shown to exhibit instability and buckling, especially when applied to models from patient imaging data [79].

The approach to estimating a corresponding pre-stress field for the diastolic femoral arterial configuration in the FSI model is following the pre-stress pipeline recently developed by Votta *et al*. for ABAQUS [80] that we now adapt for use in ANSYS Mechanical. This approach pre-stresses the initial configuration Ω_0,_ which is assumed to have been measured at diastolic pressure, by applying diastolic pressure P_diast_ and then mapping the resulting stress field σ_diast_ = σ_pre_ back onto Ω_0_ to create a pre-stressed configuration that will show no significant deformation when P_diast_ is applied again. While this approach of directly pressurising to full P_diast_ works well with linearly elastic models as demonstrated by Votta *et al*., it leads to non-physiologically high stresses in hyperelastic models due to the exponential stress-strain behaviour of hyperelastic materials. For this reason, the approach was further modified for hyperelastic models following Caimi *et al*. [81] by dividing the pressurisation stage into N consecutive steps while increasing the pressure by the same pressure increment ΔP = P_diast_/N and applying pre-stress iteratively in each step, as outlined in Figure 2. This approach was first published in 2020 for FEA simulations using Abaqus. P_diast_ for the arterial geometries were retrieved from the pulse wave database (PWD) [54] and N = 10 was chosen following Caimi *et al*. [81].

**Figure 2:**
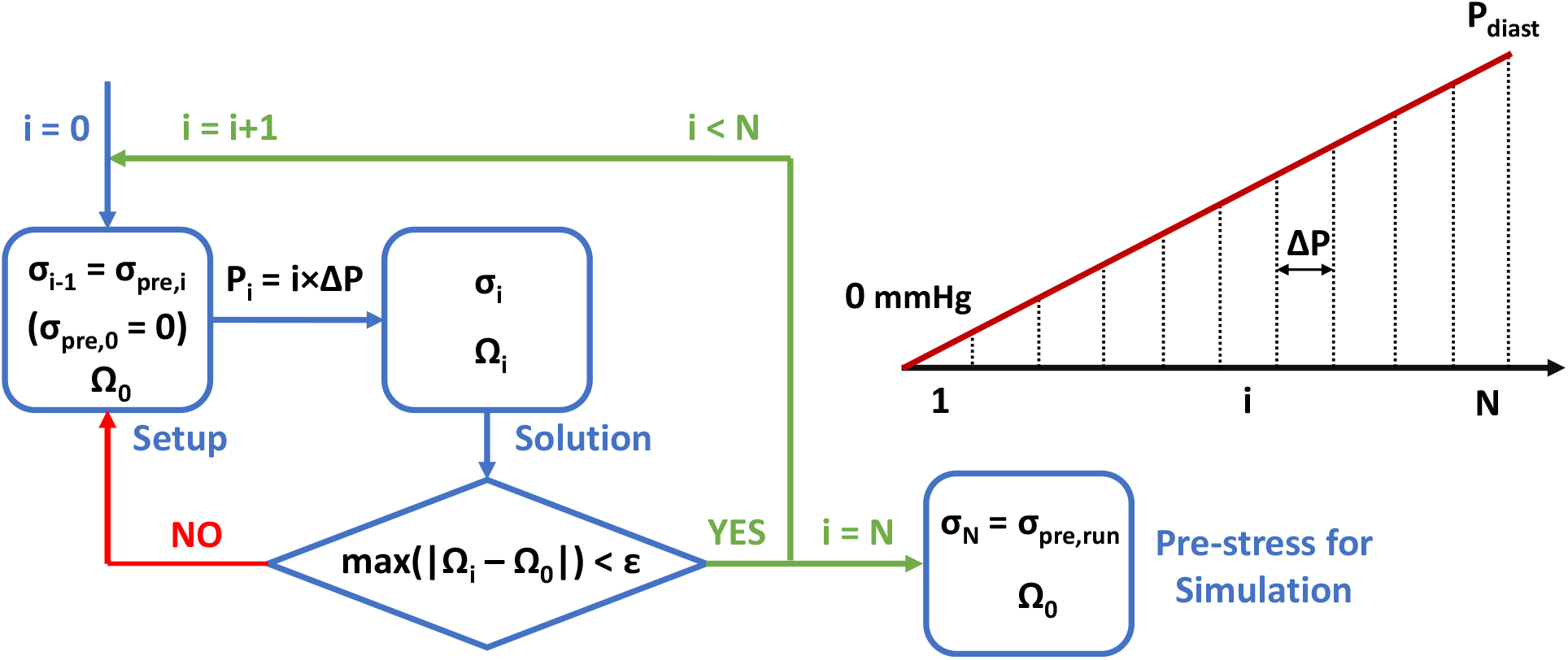
Schematic of the pre-stress pipeline for numerical simulations where the stress fields for iteratively higher pressures are calculated and the pressurised numerical model Ω_i_ is compared with a known arterial geometry Ω_0_. The linear pressure increase ΔP per pressurisation step i is determined by the diastolic arterial pressure P_diast_ and the total number of pressurisation steps N.

With the pressure in each pressurisation step i set to P_i_ = i×ΔP, pre-stress for each step is generated as follows:

1. **Initial State:** The model is pre-stressed by mapping the stress field of the previous pressurisation step σ_i-1_. This can be defined as the vector principal stresses onto the initial configuration Ω_0_. For i = 0, the configuration is in a zero-stress state.
2. **Time Step 1**: Pressurisation with P_i-1_ for 0.05 s.
3. **Time Step 2**: Linear pressure ramp-up to P_i_ over 0.05 s.
4. **Time Step 3**: Pressure is kept at P_i_ for 0.2 s to allow the stress σ_i_ and configuration Ω_i_ to stabilise.
5. **Check**: If the pressurised geometrical configuration Ω_i_ is equivalent to the initial one (max(|Ω_i_ – Ω_0_|) < ε), the pressurisation step ends. Otherwise, the step is iterated using σ_pre,i_ = σ_i_.
6. **Export**: At the end of the pressurisation step, the stress field σ_i_ is exported to serve as pre-stress σ_pre,i+1_ for the next pressurisation step.

This process is repeated until i = N = 10 and the resulting stress field is used as the pre-stress for the simulation (σ_10_ = σ_pre,sim_). Similar to Caimi et al., ε is defined as the spatial resolution of current CT scanners since patient-specific geometries will be modelled using CT scanned images in future patient-specific models. Accordingly, ε was set to 0.5 mm [82], but due to the high total number of pressurisation steps N and axisymmetric nature of the idealised geometry, a higher dimensional accuracy with max(|Ω_i_ – Ω_0_|) < 0.05 mm was achieved for every pressurisation step i.

#### 2.2.2 Fluid Model

In this study, the Carreau-Yasuda model was used to describe blood viscosity [83, 84]. The apparent viscosity η is given by the shear rate 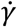, empirically determined constant parameters a, n, and λ, and viscosities 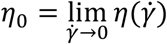 and 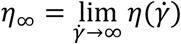

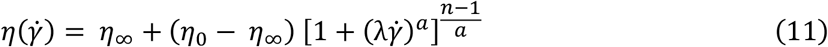

with *η*_∞_ = 0.0035 Pa·s, *η*_0_ = 0.1600 Pa·s, λ = 8.2 s, *a* = 0.64, and *n* = 0.2128 [83].

A time-dependent velocity profile was prescribed at the inlet of the fluid model as the inflow boundary condition. A matching time-dependent pressure profile was prescribed at the outlet. Boundary conditions were assumed to be uniform over the cross section of the inlet and outlet. Both the inflow and outflow velocity or pressure boundary conditions were described as eighth-order Fourier series fit:

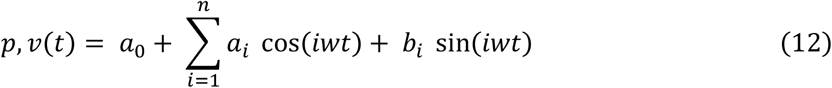

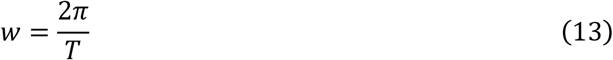

where *a*_0_ models a constant intercept term in the data and is associated with the i = 0 cosine term, *w* is the fundamental frequency of the signal, *T* is the signal’s period i.e., the cardiac cycle length, and *n* (= 8) is the number of terms in the series. The Fourier series were fitted to pressure and velocity data taken from the Pulse Wave Database (PWD) [54]. The model was fitted for two heartbeats to ensure periodicity, using the MATLAB (MathWorks, Massachusetts) Curve Fitting tool. To facilitate a nodal comparison in haemodynamic parameters between the two-way and one-way FSI models, the same fluid mesh was utilised for all simulation of a given age or wall thickness.

The model was run as a transient analysis with a Second Order Backward Euler transient scheme with a time step size of 10^−3^ s and high resolution (2nd order iteratively bounded) advection scheme. While the timestep sensitivity analysis allowed for a larger timestep, this step size was chosen to allow for direct comparison of CFX results with patient data from the PWD which provides data points every two milliseconds. A timestep was considered as converged when the Root Mean Square (RMS) residuals of the mass conservation equation and the momentum conservation equations were all below a residual target of 10^−4^. The model was run for three cardiac cycles for the studied patients and only the last cycle was analysed to eliminate transient start-up effects from the results.

During result analysis, the Womersley number *α* was calculated, which is a dimensionless parameter that describes the relationship between the viscosity and pulse flow frequency in a pulsating internal pipe flow. It has been demonstrated that when *α* < 1, the flow is expected to follow the pulsating pressure gradient faithfully, and the velocity profiles exhibit a parabolic shape. When *α* > 1, the velocity profiles are no longer parabolic, and the flow is out of phase in time with respect to the pressure gradient [85]. *α* > 1 are commonly observed in blood vessels with a diameter larger than 3 mm such as the femoral arteries [86, 87].

The average dynamic viscosity *η*_avg_ for the Carreau-Yasuda blood model is calculated with the average shear rate defined as:

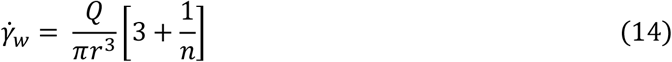

where Q is the average volumetric flow rate per cardiac cycle, *r* is the average internal radius and *n* is the power law index. The Womersley number can then be calculated as [88]:

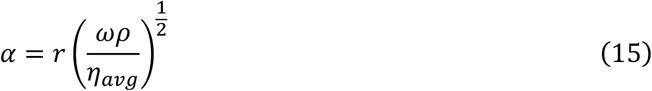

with fluid density *ρ* and angular frequency *ω* derived from the cardiac cycle length (period) [89]

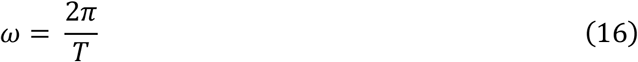

The numerical model simulating the arterial phantoms uses a Newtonian water model.

### 2.2.3 System Coupling

ANSYS System Coupling is a computational framework which allows individual physics solvers to communicate with one another. During execution, one- or two-way data transfers between a source region and a target region are performed for coupling participants [90]. In our system, data transfer regions are defined at the outer surface of the fluid volume and on the luminal surface of the structural model.

The workflow of the strong two-way coupling algorithm for FSI modelling is shown in Figure 3. After convergence of the fluid solver, forces are transferred to the structural solver and interpolated to the structural mesh. A solution from the structural solver is obtained with those fluid forces as boundary conditions resulting in a deformation of the structural mesh. These displacement values are transferred back to the fluid solver and interpolated to the fluid mesh which results in deformation of the fluid domain. This process is repeated for each time step until both the individual solvers and the coupling data transfers have met an RMS convergence target of 10^−2^ [14, 91]. Fluid mesh deformation was calculated using a displacement diffusion model without remeshing. The flow solution was stabilised to prevent divergence by setting a velocity-dependent source coefficient in the pressure equation at the arterial wall and monitoring monotonical convergence for a given time step by plotting the forces and displacements at the arterial wall boundary [92].

**Figure 3:**
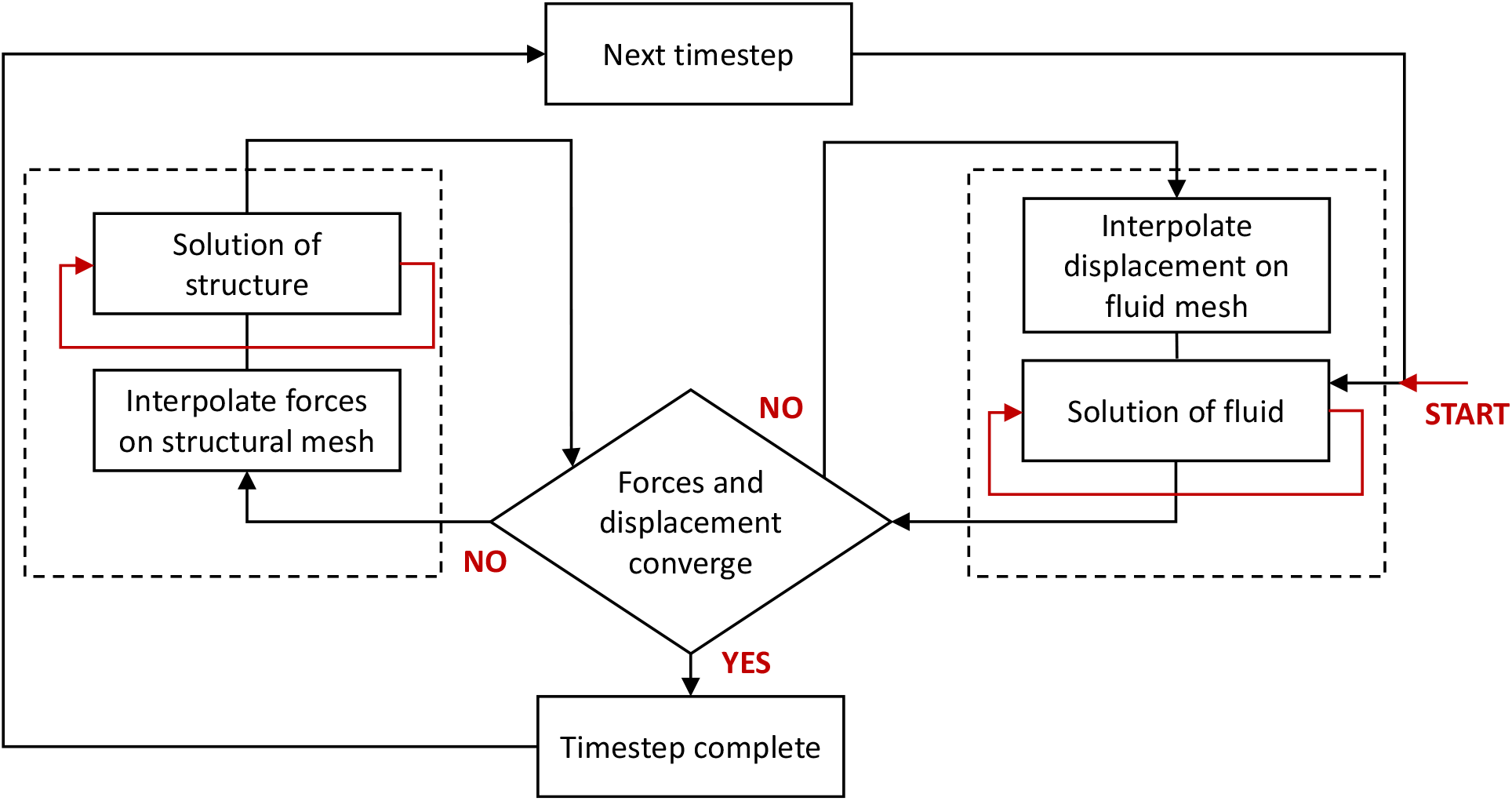
Fluid-structure interaction solver pipeline using strong two-way coupling. Convergence for each timestep requires convergence of the individual solvers (ANSYS CFX and Mechanical) and convergence of the data transfers.

Results from two-way coupling were compared with weak one-way coupling, which only includes the force transfer from the fluid to the structural solver and assumes the effect of the structural mesh displacement on the fluid solution to be negligible.

## 3 Results

### 3.1 Coupled *in silico* and *in vitro* Models of Arterial Phantoms

Manufacturing of six sample groups of arterial phantoms was successfully undertaken by 3D FDM printing of moulds and casting of PlatSil Gel-0020, Sylgard 184 PDMS, and Smooth-On Solaris silicones with wall thicknesses of 0.8 mm, 1.2 mm, and 1.6 mm. For all three materials and wall thicknesses (n = 5 per group), the final manufactured phantom wall thickness was measured to be approximately 10% larger than the respective wall thickness of the negative mould design for all materials (Figure 4A), likely due to the swelling properties of the PVA mould material. This effect needs to be considered when tuning a model for desired mechanical properties because mechanical compliance is directly related to wall thickness.

**Figure 4:**
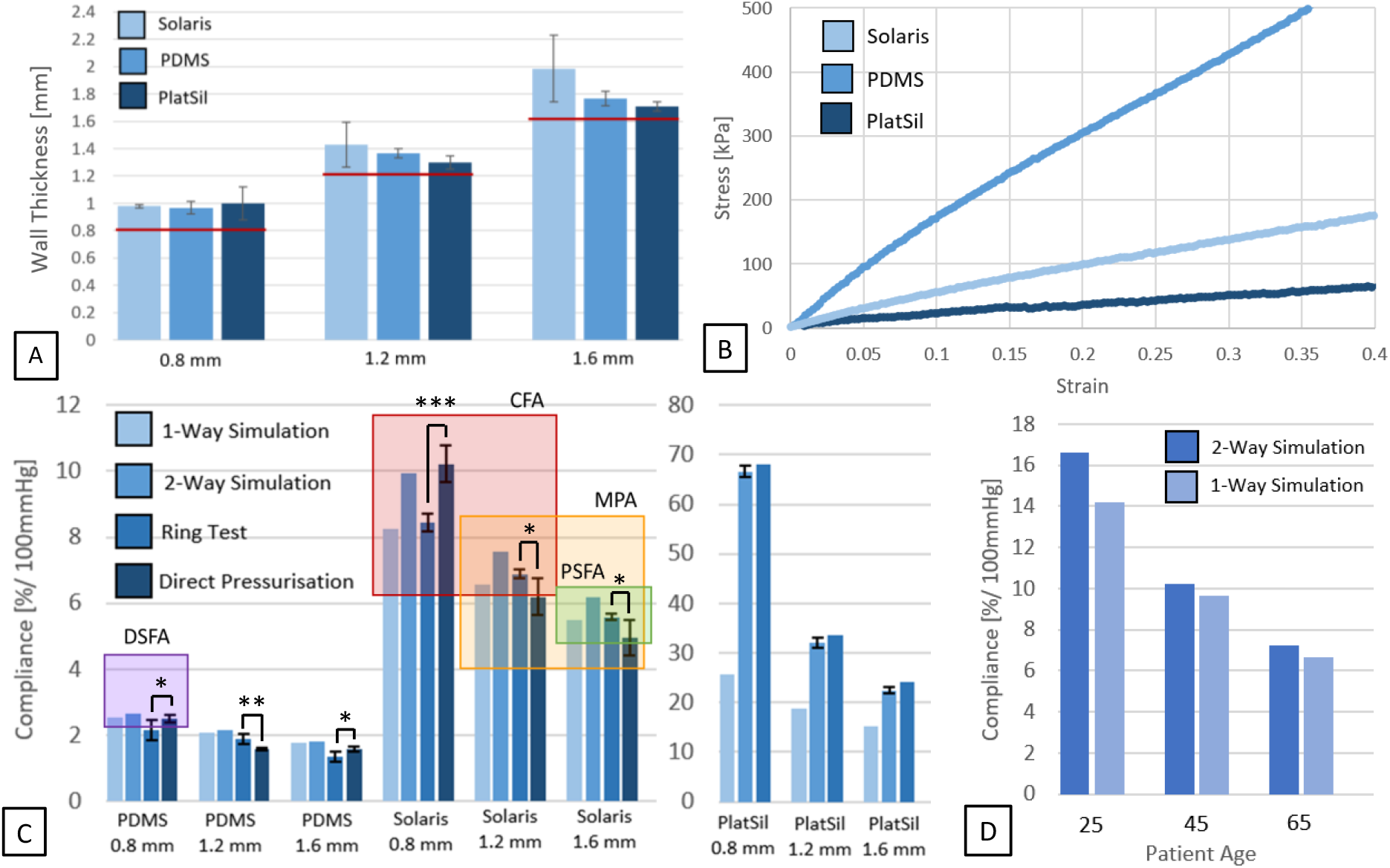
Experimental and numerical results of silicone arterial phantoms and arterial patient models. A: Tensile behaviour of PlatSil Gel-0020, Sylgard 184 PDMS, and Smooth-On Solaris as illustrated by exemplary samples; B: Measured (n=5) wall thicknesses of PlatSil Gel-0020, Sylgard 184 PDMS, and Smooth-On Solaris samples vs. wall thickness in CAD mould model (red line); C: Circumferential compliance of arterial phantoms as calculated from direct pressurisation (120/80) measurements (n=5), ring test data (n=5), and one-way and two-way FSI simulations with corrected measured wall thickness (4A). Comparison with CFA: Common Femoral Artery, MPA: Midgenicular Popliteal Artery, PSFA: Proximal Superficial Femoral Artery, DSFA: Distal Superficial Femoral Artery. Boxes represent typical compliance ranges for those regions in patients with and without PVD [93]. D: Circumferential compliance of numerical patient models for different ages. Significance testing compliance measurements from ring tensile testing versus direct pressurisation is denoted by *: 0.01<p<0.05, **: 0.001<p<0.01, ***: p<0.001.

The Young’s modulus of the arterial phantoms was mechanically examined by direct pressurisation (n = 5 per group) and tensile ring testing (n = 5 per group) and compared with one-way and two-way FSI simulations informed by the measured phantom thicknesses (Figure 4A) and stress-strain measurements (Figure 4B). All ring samples broke in a straight line along the axial direction of the sample. The stress-strain behaviour of the tested silicones is in agreement with published literature [94] and representative stress-strain curves encountered during ring tensile testing are illustrated in Figure 4B. No compliance data under direct pressurisation could be generated with PlatSil samples due to their large deformation. A two-tailed unpaired t-test showed differences between the measured compliance values of the direct pressurisation and the ring testing method that are considered at least moderately statistically significant (p <0.05) in all sample groups, with the most significant difference measured for the 0.8 mm thick Solaris samples (p = 0.0002), which is the most compliant test group, where both mechanical testing methods were successfully applied. Overall, the difference between the testing method ranged from 10 - 20% for each sample group.

For all tested wall thicknesses, the compliance of PlatSil silicone samples were greater than *in vivo* compliance measurements of peripheral arteries described in the literature (Figure 4C) and the numerical femoral arterial models (Figure 4D). Conversely, the compliance of PDMS samples was lower than the compliance measured in all numerical patient models and most *in vivo* patient measurements described in literature apart from the distal superficial femoral artery in older patients with PVD [95]. Smooth-On Solaris sample matched the compliance range of the numerical and *in vivo* patient models most closely out of the tested silicone materials (Figure 4D). For matching wall thicknesses of 1.2 mm, the two-way FSI Smooth-On Solaris model best mimicked the compliance behaviour of the 65-year-old two-way FSI patient model (7.559%/100mmHg vs. 7.237%/100mmHg). In both silicone and patient models where the fluid mesh deformation is neglected (one-way coupling), the measured compliance is reduced compared to a two-way coupled model. This effect depends on the compliance of the structural model and consequently on the health and age of the patient model with a reduction in compliance from 16.635 %/100mmHg to 14.188%/100mmHg (14.71%), from 10.253%/100mmHg to 9.654%/100mmHg (5.84%), and from 7.237%/100mmHg to 6.651%/100mmHg (8.11%), for a 25-year-old, 45-year-old, and 65-year-old patient model respectively. While the arterial wall material model of the 65-year-old patient is stiffer compared to the 45-year-old patient, the increase in systolic blood pressure in the older patient model leads to a larger percentage difference in between one-way and two-way FSI modelling in this case.

### 3.2 One-Way and Two-Way Fluid-Structure Interactions in Small-Diameter Blood Vessels

The arterial structural models were assessed regarding internal diameter changes along the cardiac cycle (*D*_*i,mod*_) compared with measurements taken from PWD (*D*_*i,pat*_) (Figure 5A) to confirm a physiological deformation behaviour for each patient age. The resulting curves were compared using the normalised Euclidean dissimilarity metric or normalised Euclidean distance [96, 97], which is defined as:

**Figure 5:**
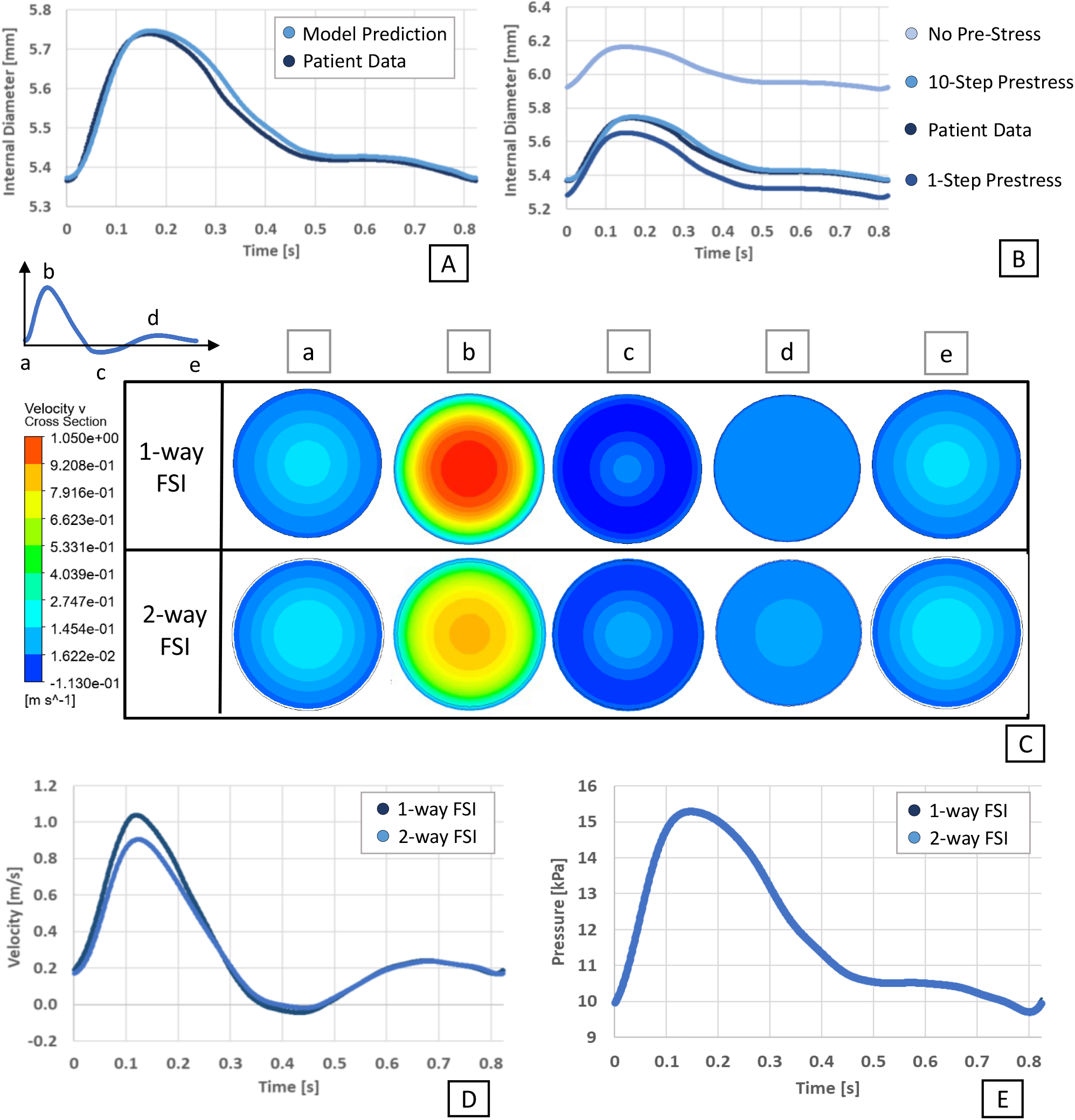
Impacts of pre-stress and fluid-structure interactions (FSI) on arterial fluid dynamics in a 25-year-old patient model. A) Artery internal diameter change during the cardiac cycle with patient data from [54]; B) Arterial wall deformation for different pre-stress states; C) Velocity profiles for one-way and two-way FSI along cardiac cycle; D) Central axial values for velocity along cardiac cycle for one-way and two-way FSI; E) Central axial values for pressure along cardiac cycle for one-way and two-way FSI.

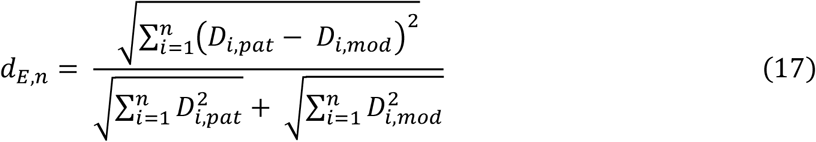

Values of d_E,n_ fall in the interval [0,1]. A dissimilarity value of 0 indicates that the two curves are identical and the larger the value of the dissimilarity coefficient the bigger the difference between the two data sets. The average difference between patient compliance data and the model estimation along the cardiac cycle is d_E,n_ < 1.6×10^−3^ for all age models, indicating a high degree of similarity (Figure 5A).

A comparison of patient data and two-way FSI results with a ten-step iterative pressurisation pipeline to a model with no pre-stress shows that the diameter of the artery would be overestimated (d_E,n_ = 4.4×10^−2^) throughout the cardiac cycle while the difference between systolic and diastolic diameter would be underestimated (Figure 5B). This is because the intramural stresses of the arterial wall captured *in vivo* by medical imaging were neglected. Assuming a zero-stress state of the diastolic geometry leads to the hyperelastic material models showing a stiffer behaviour at the resulting higher deformations at systole. Conversely, a model where the pre-stress has been calculated by simply applying the full diastolic pressure in one step is not appropriate (Figure 5B) either, as the hyperelastic material nonlinearly responds with elevated wall stresses to the large deformation of the model. The final nodal pre-stresses of the one-step pressurisation pipeline were approximately 20% higher than those generated with the iterative ten-step pipeline, leading to a contraction of the model below diastolic dimensions (d_E,n_ = 8.7×10^−3^) when diastolic pressure was applied again via two-way FSI coupling.

The generated velocity profiles for different key times of the cardiac cycle are in good agreement with literature [98]. The effect of the Womersley flow on the velocity profile can be observed for both one-way and two-way FSI results (Figure 5C) with a reduction in peak velocities in the two-way FSI model. The average Womersley number of 4.06 for this model is in good agreement with literature on femoral arteries [86, 87]. In two-way FSI, the elastic effect of the deforming arterial wall reduces the temporal maximum and minimum velocity values (Figure 5D) for a given pressure profile (Figure 5E).

In Figure 6, a node-wise comparison was performed in the relaxation phase of the cardiac cycle at 0.271 s, where central axial velocity values are equal between one-way and two-way FSI results (as shown in Figure 5D). Figure 6’s node-wise comparison shows differences between local velocity, pressure, deformation, and wall (intramural) stress values. In this study we calculated the Pearson’s product-moment correlation coefficient (R^2^) of the one-way and two-way data for the velocity and pressure parameters of the fluid model and the radial deformation and wall stress of the structural model. An R^2^ value greater than 0.7 indicates a high positive correlation [99].

**Figure 6:**
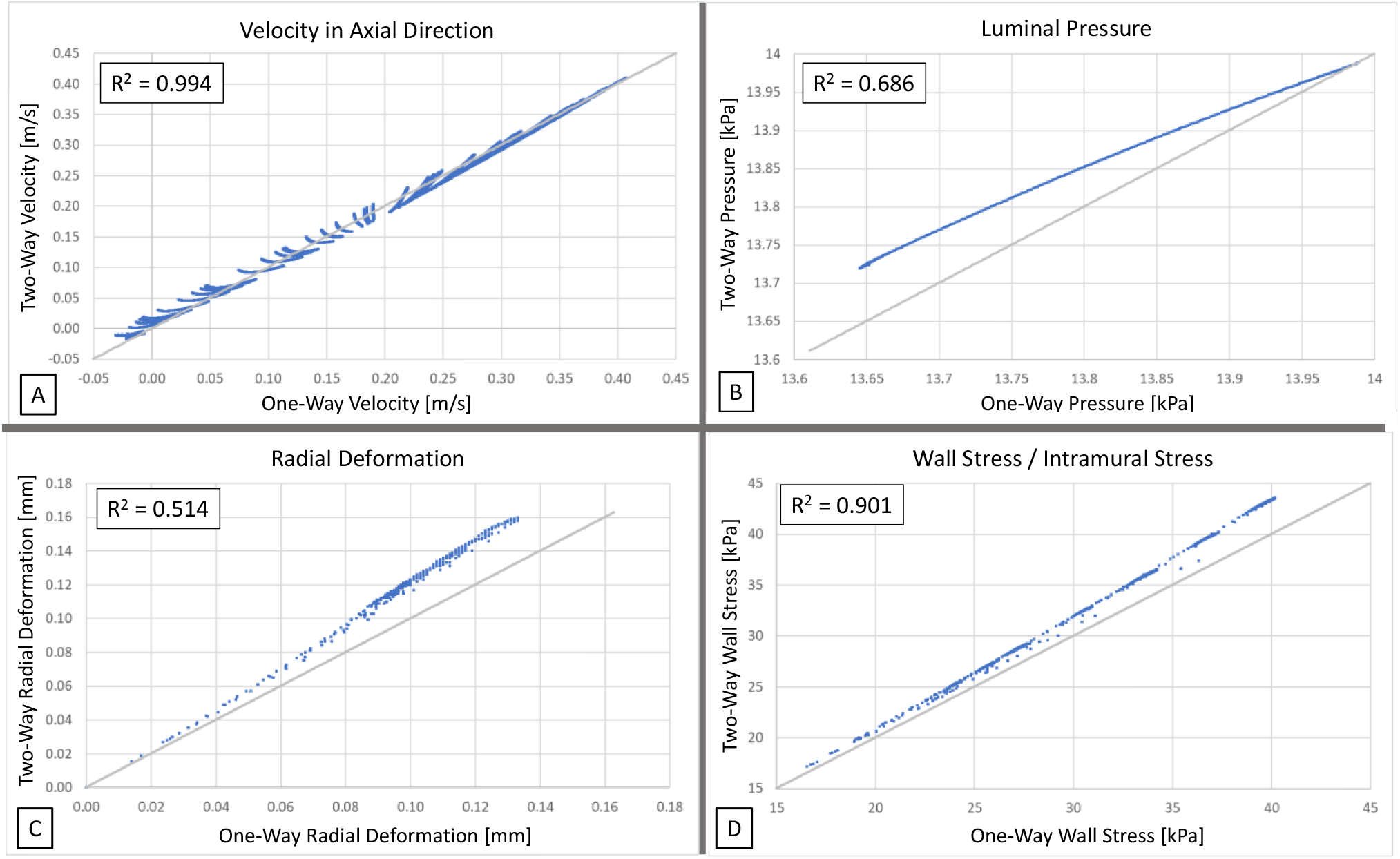
Node-wise comparison of one-way and two-way FSI results at 0.271 s in the cardiac cycle of the 25-year-old patient model. This average of these nodes represents an identical average velocity of 0.41 m/s for both one-way and two-way FSI simulations (shown in Figure 5D). Node-wise differences between two-way FSI (y-axes) and one-way FSI (x-axes) in A) velocity in an axial direction, B) luminal pressure, C) radial deformation, and D) intramural stress (von Mises stress) are calculated, superimposed with a grey diagonal line and an R-correlation to describe agreement between two-way and one-way FSI simulations. The grey line indicates hypothetical data points with the same prediction of 1-way and 2-way FSI for a given node.

For a timepoint where both models have the same central axial velocity value, the spatial differences in velocity due to the deforming fluid mesh are small (R^2^ = 0.994; Figure 6A). While the temporal similarity between the one-way and two-way pressure profiles is high (Figure 5E), the spatial correlation between the pressure profiles is low (R^2^ = 0.686; Figure 6B) for a node-wise comparison and local pressure values are underestimated for one-way coupling except for central nodes. The nodal differences for deformation (R^2^ = 0.514; Figure 6C) are high, which is in agreement with the reduction in compliance for one-way FSI models described in Figure 4D. In the structural model, WS profiles have a high spatial correlation (R^2^ = 0.901; Figure 6D) but are underestimated at all nodes. This difference is most pronounced for the maximum wall stresses which occur at the luminal arterial wall model (8.05%). Although one-way and two-way FSI exhibit smaller spatiotemporal differences in velocity profiles, more substantial differences were found in parameters influencing arterial compliance.

## 4 Discussions

A relatively small subset of computational fluid dynamics (CFD) publications considers the impact of fluid-structure interactions (FSI) when modelling the flow of blood through highly compliant peripheral arteries [100]. These mechanical interactions between flowing blood and the artery wall at different artery locations, stages of the cardiac waveform, and disease progression, are central toward understanding and diagnosing cardiovascular pathology, predicting its future trajectory, providing relevant surgical training models and simulations, and designing new cardiovascular interventions.

In this study, one-way and two-way FSI models were analysed for the simulation of patient cases of different ages. Two-way FSI models are more computationally expensive than one-way FSI models. For instance, staggered two-way FSI simulations of problems with comparable mass densities such as blood flow-artery interactions create numerical instabilities [101], especially in cases where the structure is highly deformable. These challenges associated with two-way FSI coupling can become especially significant in simulations with many sub-steps or simulations that involve complex geometries such as patient-specific vascular models. The choice between one-way and two-way FSI simulations depends on the mechanical compliance of the blood vessels, which is influenced by factors such as age and health status. For most compliant case analysed in this study, a 25-year-old patient without PAD, a compliance reduction of 14.71% from the two-way FSI model to the one-way FSI model was measured.

The acceptability of the reduction of compliance prediction resulting from neglecting the deformation of the fluid mesh needs to be considered for the specific case to be simulated. The smaller the model compliance and the larger the parameter differences to be measured between models, the more acceptable the use of one-way FSI. For vascular applications, the differences in average mechanical compliance between peripheral arteries of the lower limbs with and without PAD ranges from 13.6 - 56.0% depending on location [95] with a reduction of 38.7% [102] and 47.0% [103] for the femoral artery specifically. The difference between one-way and two-way FSI predictions might be unacceptable in applications where small changes in compliance are considered significant such as studies where vascular compliance is utilised as a functional diagnostic parameter, e.g. to diagnose stenosis due to smooth muscle cell proliferation (2.5%) [104]. Other examples are studies on the effect of treatments on arterial compliance such as ultrafiltration in haemodialysis (4.8%) [105], or the effect of excess bodyweight (12.3%) [106], medication and dietary supplements (12 - 15%) [107, 108].

Since one-way FSI does not feed information on arterial wall deformation back into the blood flow calculations, a loss of accuracy is expected when determining haemodynamic parameters. While temporal differences in the predicted velocity profiles across the cardiac cycle were pronounced with an overestimation in peak velocity in the one-way coupled model, for a timepoint where both models have the same central axial velocity value, the spatial differences in velocity due to the deforming fluid mesh are small. The one-way FSI approach led to an underestimation of the maximum von Mises stress within the structural model especially around the vessel lumen, with high intramural stresses having been indicated as a risk factor in intimal hyperplasia since they can trigger both endothelial damage and smooth muscle cell proliferation [109]. The most pronounced nodal differences were noted for local WS and deformation, which were underestimated in one-way FSI. These local differences may be non-negligible in studies where local arterial wall distensions and stress concentrations are studied such as aneurysm models [110]. For the numerical modelling of the arterial wall, the importance of applying an appropriate pre-stress to the structural model is widely established in the literature [111]. Our study confirmed that neglecting arterial wall pre-stress and assuming a zero-stress state at diastole leads to an non-physiological overestimation of the arterial diameter throughout the cardiac cycle while simultaneously reducing the distensibility of the structural model between systole and diastole (Figure 5B). Additionally, comparing the deformation behaviour of arterial models pre-stressed in one pressurisation step to those informed by an iterative ten-step pipeline, an iterative pre-stress process may generate a more accurate diastolic pressurised geometrical configuration for hyperelastic arterial wall models.

During the mechanical characterisation of the femoral arterial phantoms, compliance measurements of the experimental models utilising direct pressurisation and ring testing have been compared. While direct pressurisation can potentially replicate the physiological deformation of an artery more closely, this testing rig may not be easily available since it requires a laser micrometer, a pump system which can replicate physiological pressures and frequencies, and a coupled pressure gauge. Ring testing has been described in literature as an alternative to direct pressurisation for tissue-engineered vascular substitutes [53], and has produced acceptable results in this study for compliance testing of silicones with mean differences of 10 - 20% between the two methods. The differences between the measured compliance values of the direct pressurisation and the ring testing method were at least moderately statistically significant in all sample groups, with the most significant difference measured in the most compliant test group. These differences may be partially due to the cyclic and fast deformation during direct pressurisation which affects the distensibility of silicone materials [112]. Overall, ring tensile testing may provide a useful alternative to direct pressurisation where no pressurisation rig is available, the sample length or number is insufficient, or the sample deformation exceeds the micrometre measurement area.

For all tested wall thicknesses, the compliance of PlatSil silicone samples ranged above compliance measurements of peripheral arteries described in the literature while the compliance of PDMS samples was lower than the compliance measured in healthy *in vivo* patient measurements [95]. Smooth-On Solaris sample matched the compliance range of the numerical and *in vivo* patient models most closely out of the tested silicone materials. In arterial phantom manufacturing, PDMS is currently a commonly utilised material [113] in both stiff models, which are fabricated as a large block of silicone with a cavity replicating the lumen of the patient geometry, and in compliant models, which are fabricated using an offset from the lumen geometry with variable thickness. While PDMS models are regularly described as compliant, the specific stiffness or compliance of the model is often not quantified [114-116]. PDMS phantoms wall thicknesses of 1 - 3 mm, which replicate the arterial wall thickness of commonly studied arteries from aorta to peripheral vessels and are utilised in commercial models [117], are too stiff to replicate physiological arterial deformation. The stiffness of PDMS may be lowered by reducing the crosslinking ratio in the silicone gel mix [118], however, the shape stability and tensile strength of the model is reduced in off-ratio mixtures [119, 120] and residual monomers are free to diffuse within the bulk material and leach from the model into any adjacent testing fluids or cell culture systems [121]. For PDMS, a closer compliance match for the same wall thickness is expected for the simulation of stiffer peripheral arteries in the case of older patients or patients with vascular disease. Compliance could also be tuned by changing the wall thickness of the arterial phantoms until sufficient similarity is achieved, although non-physiological wall thicknesses may be required [122].

Depending on the application of the experimental vascular model, requirements regarding transparency, compliance, and personalisation vary. While an opaque compliance-matched model may be suitable for surgical training, many experimental flow measurement techniques such as particle image velocimetry (PIV) require a high level of material transparency [123]. The direct additive manufacturing of transparent compliant arterial phantoms remains challenging due to a lack of suitable materials. While some manufacturers such as Formlabs (Somerville) and 3DSystems (Rock Hill), have developed flexible and translucent resins and many other companies are distributing clear thermoplastic polyurethane (TPU) filament for FDM printing, there is currently no material commercially available which combines the requirements for transparency, elasticity, and compliance to facilitate direct fabrication of arterial phantoms with physiological compliance. While the additive manufacturing of transparent elastic silicone gels has been demonstrated in the literature [124] and companies offering silicone AM services such Spectroblast AG (Zürich) are emerging, the suitability of these technologies for arterial phantom manufacturing regarding printing resolution, scalability, and material properties has yet to be demonstrated. For this reason, silicone casting via a 3D printed mould remains a commonly used method for vascular model fabrication [113].

Several limitations remain in this new experimental-numerical approach for the development of physiologically complaint arterial phantoms. All the simulations of the present study are based on a flat profile boundary condition, which does not replicate physiological flow. However, sufficient entry and exit length has been granted for the Womersley flow profile to fully develop in good agreement with published literature. Furthermore, the arterial wall in our present study is regarded as a single layer with constant thickness. In patient-specific cases, the distribution of vascular wall thickness may be different along the luminal wall of the artery. This simplification has been applied because layer-specific geometric data are seldom collected in clinic and cannot usually be extracted from MRI or CTA scans. In the experimental model, the pump utilised in this study generated systolic and diastolic pressures in a sinusoidal waveform and could not be programmed to replicate the exact pressure waveform within the femoral artery. Since only the systolic and diastolic pressure and deformation values are utilised for the calculation of circumferential compliance, this limitation is expected to have had little effect on the measured compliance values.

In future work, the described additive manufacturing and moulding methodology may be utilised to create more complex and patient-specific models. While the biocompatibility of PlatSil [125] and PDMS [126] have been demonstrated in the literature, Solaris has been given little academic attention as a substrate for bioengineered models and might be an interesting candidate for cellularised vascular models or actuators to mimic vascular biomechanics due to its physiological compliance range and transparency. Solaris’s optical properties might also be useful for the experimental analysis of fluid flows e.g. via PIV, where PDMS is already established as a common material for silicone vascular models [113].

## 5 Conclusion

In this paper, a numerical simulation of an idealised femoral artery is performed via one-way and two-way FSI simulations and compared with experimental data collected from arterial phantoms fabricated *via* additive manufacturing. For the most compliant case analysed in this study, a 25-year-old patient without PAD, a compliance reduction of 14.71% from the two-way FSI model to the one-way FSI model was measured with the largest spatial differences for a given timepoint noted for nodal WS and deformation results, which were underestimated in one-way FSI. Temporal differences in the predicted velocity profiles across the cardiac cycle were pronounced with an overestimation in peak velocities for one-way coupling. These local differences may be significant in studies where small to medium differences in compliance or local quantitative haemodynamic and structural values are of interest. Despite the increase in computational cost, two-way FSI simulation should be considered for small-diameter vascular applications unless it can be demonstrated that the effect of the deforming fluid mesh is negligible for the simulated model. Additionally, comparing the deformation behavior of arterial structural models pre-stressed in one pressurisation step to those informed by an iterative ten-step pipeline, an iterative pre-stress process may generate a more accurate diastolic pressurised geometrical configuration for hyperelastic arterial wall models.

Additive manufacturing and silicone-casting facilitate the fabrication of arterial phantoms with physiological compliance for physiological wall thicknesses and allow for personalisation of models using patient data. During the mechanical characterisation of the femoral arterial phantoms, compliance measurements of the experimental models utilising direct pressurisation and ring testing have been compared and showed moderate differences of 10 - 20% between the two methods. Ring tensile testing may provide a useful alternative to direct pressurisation for measuring vascular model compliance where direct pressurisation is not available or applicable. For all tested wall thicknesses, the compliance of PlatSil silicone samples ranged above compliance measurements of peripheral arteries described in the literature while the compliance of PDMS samples was lower than the compliance measured in healthy *in vivo* patient measurements. Smooth-On Solaris sample matched the compliance range of the numerical and *in vivo* patient models most closely out of the tested silicone materials.

Arterial phantom modelling offers future use in vascular disease research, implant development, and surgical training where physiological arterial wall behaviour is essential. Compliant vascular models are useful because they can simulate the mechanical behaviour of blood vessels in the human body, allowing researchers and clinicians to study and understand complex cardiovascular diseases and conditions. These models can replicate the physical properties of the human vasculature, including its elasticity, compliance, and resistance to blood flow, and can be used to evaluate the performance of medical devices and treatments, such as stents and balloons, before they are used on patients. Compliant vascular models also provide a valuable tool for education and training, allowing medical students and professionals to practice complex procedures and develop their skills in a safe and controlled environment. Ultimately, the use of compliant vascular models can lead to improved patient outcomes and better healthcare for individuals with cardiovascular disease.

## Declaration of Competing Interest

The authors declare that they have no known competing financial interests or personal relationships that could have appeared to influence the work reported in this paper.

## Acknowledgments

This work was supported by a GOstralia!-GOzealand! GmbH Scholarship to SS, a Bionics Queensland Grand Challenge Award to SS and MCA, an Advance Queensland Fellowship to MCA (AQIRF1312018), an Australian Research Council DECRA Fellowship to MCA (DE220100757), and a British Heart Foundation Imperial Centre of Research Excellence Award (RE/18/4/34215) to SP.

## Notes

### Competing Interest Statement

The authors have declared no competing interest.

